# Bridging the Gap in Immunotherapy Prediction: The AGAE Score as a Pan-Cancer Biomarker for Immune Checkpoint Inhibitor Response

**DOI:** 10.1101/2024.10.15.618438

**Authors:** Bicheng Ye, Jun Fan, Qingren Meng, Qingxiao Liang, Aimin Jiang, Pengpeng Zhang

**Author notes:** These authors have contributed equally to this work and share first authorship (Bicheng Ye, Jun Fan, Qingren Meng). **Correspondence:** Pengpeng Zhang; Aimin Jiang; Xiaoqing Liang.

## Abstract

**Background:** Immune checkpoint inhibitor (ICI) therapy efficacy varies among cancer patients, necessitating precise predictive biomarkers for optimized treatment strategies.

**Methods:** We developed the Adaptive best subset selection algorithm and Genetic algorithm Aided Ensemble learning (AGAE) score through multi-cohort transcriptomic analysis of ICI-treated patients. The AGAE score incorporated gene-pairing, Adaptive Best Subset Selection for feature optimization, and a Genetic Algorithm for optimal basic learner identification. We explored correlations between AGAE score and immune microenvironments using multi-omics data. Potential targets were screened using 17 CRISPR datasets and validated through in vitro and in vivo experiments.

**Results:** The AGAE score demonstrated robust predictive power for ICI therapy outcomes, with lower scores correlating with enhanced treatment response. The AGAE score outperformed published signatures and conventional biomarkers. Lower AGAE scores were associated with increased immune cell infiltration, higher immunogenicity, and enhanced antitumor immune activity. The CEP55 was identified as a potential key target driving immune evasion through AGAE scoring and CRISPR screening. Experimental validation showed CEP55 downregulation attenuated tumor cell malignancy and augmented ICI therapy efficacy by modulating T cell responses.

**Conclusions:** The AGAE score was a potent predictor of ICI therapy efficacy, facilitating refined patient stratification. CEP55’s role in the tumor microenvironment’s immune response highlights its potential as a therapeutic target. Targeted interventions against CEP55 may improve immunotherapy precision.

## Introduction

Immune checkpoint inhibitors (ICIs) have revolutionized cancer treatment, significantly improving survival outcomes for many patients^1–3^. The utilization of ICIs among eligible cancer patients in the United States has grown exponentially, from 1.54% in 2011 to 43.63% by 2018^4^. However, their effectiveness remains suboptimal, with only 12.46% of patients showing favorable responses^4^. The efficacy of ICIs varies considerably across different cancer types and individual cases^5^. Researchers have identified several indicators associated with inflamed tumors that may predict ICI response and prognosis, including tumor-infiltrating lymphocytes (TILs), tumor mutation burden (TMB), Immunoscore, and Gut OncoMicrobiome Signatures. However, these biomarkers have demonstrated limited predictive capabilities^6–9^. Currently, no universally applicable biomarker can accurately forecast ICI efficacy across various cancer types, hindering the optimization of patient selection and the development of effective combination therapy strategies.

The Tumor Immune Microenvironment (TIME), consisting of various immune cells, significantly influences cancer outcomes and ICI efficacy^10,11^. Understanding the relationship between TIME characteristics and ICI outcomes could improve patient selection and enhance cancer immunotherapy effectiveness. However, the intricate immune cell interactions often lead to conflicting interpretations, necessitating more sophisticated analytical approaches. Conventional methods like immunohistochemistry (IHC) and flow cytometry, commonly used to examine TIME, have limitations in simultaneously evaluating multiple cellular markers. Single-Cell RNA Sequencing (scRNA-seq) offers high-resolution transcriptomic analysis but faces practical challenges in clinical contexts due to financial constraints and stringent sample quality requirements^12 13^. In contrast, Bulk RNA sequencing (bulk RNA-seq) demonstrates remarkable adaptability with suboptimal samples and provides comprehensive transcriptome profiles from heterogeneous tissue specimens. Thus, exploring immune-related genes at the bulk RNA-seq level to characterize TIME is a reliable method^14^.

Ensemble learning techniques have demonstrated superior accuracy in biological predictions using transcriptomics data compared to single machine learning approaches^15–17^. However, significant challenges persist, including the risk of overfitting, handling diverse transcriptomic data distributions, and the lack of automated methods for selecting basic learners^18^. To address these issues, we developed the Adaptive best subset selection algorithm and Genetic algorithm Aided Ensemble learning score (AGAE score) for predicting ICI therapy outcomes in cancer patients. Our approach incorporates a gene-pairing strategy to minimize batch effects^19^, uses the Adaptive Best Subset Selection (ABESS) algorithm for optimal feature identification, and employs a genetic algorithm (GA) to automate basic learner selection^19 20^. This culminated in a web-based tool that calculates the AGAE score (https://ici-theaphy-gms.shinyapps.io/my_shiny_app/), offering predictive insights for pan-cancer patients undergoing ICI therapy. This novel approach aims to provide a more reliable and versatile predictor of ICI response, potentially enhancing treatment outcomes and guiding clinical decision-making in cancer immunotherapy.

## Methods

### Data Collection and Processing

We analyzed bulk transcriptomic profiles and clinical data from immunotherapy patients across 11 bulk RNA-Seq datasets. This comprehensive collection included: 4 Skin Cutaneous Melanoma (SKCM) cohorts (Łuksza SKCM^21^, Gide SKCM^22^, Riaz SKCM^23^, and Hugo SKCM^24^), 1 Renal Cell Carcinoma (RCC) dataset (Bruan RCC^25^), 1 Urothelial Carcinoma (UC) study (Mariathasan UC^26^), 1 Glioblastoma Multiforme (GBM) cohort (Zhao GBM^27^), 1 Gastric Cancer (GC) dataset (Kim GC^28^), and 2 Non-Small Cell Lung Cancer (NSCLC) cohorts (OAK NSCLC^29^ and POPLAR NSCLC^30^). The Hugo cohort^24^ encompasses 27 pre-treatment tumor specimens from 26 individuals, while the Zhao cohort^27^ includes 34 pre-treatment tumor samples derived from 17 patients. For these two datasets, we randomly chose one tumor sample per patient to ensure data independence. We combined six ICI RNA-Seq cohorts to create a large dataset (n=1031), which included RCC (n=172), UC (n=298), SKCM (n=244), and NSCLC (n=317). These six cohorts were Braun, Mariathasan, Łuksza, Gide, Riaz, and OAK. This combined cohort was then randomly divided into two subsets: a training set (70%, n=722) and a validation set (30%, n=309). The remaining five ICI RNA-Seq cohorts were merged to form an independent testing set (n=211), as detailed in Supplementary Table 1. For the transcriptome signature comparison, we employed the ComBat function from the sva R package to mitigate batch effects^31^.

A dataset, comprising normalized transcriptomic and genome data from 9,815 patients with complete survival information across 30 The Cancer Genome Atlas (TCGA) cohorts (Supplementary Table 2), was obtained from the University of California Santa Cruz (UCSC) Xena database (https://xenabrowser.net). Three cancer types were excluded from the analysis due to their inherently high immune cell content: diffuse large B cell lymphoma (DLBC), acute myeloid leukemia (LAML), and thymoma (THYM). This exclusion was based on previous research findings^32^.

We also acquired data from a previously published pan-cancer scRNA-seq study^33^. This comprehensive dataset comprised samples from various tissue types: 31 normal samples, 54 adjacent tissue samples, and 148 tumor samples. For subsequent investigation, 829,521 cells, chosen after rigorous selection, were subjected to the Seurat R package^34^. Gene expression data were normalized using the LogNormalize method with a scale factor of 10,000. We then identified the 2,000 most variably expressed genes and scaled them prior to performing principal component analysis (PCA). To address batch effects, we utilized the Harmony R package^35^. The entire analysis pipeline was executed with precision using both the Harmony and Seurat packages. This process encompassed several key steps: NormalizeData, FindVariableFeatures, ScaleData, RunPCA, FindNeighbors, FindClusters, and RunUMAP. Additionally, we obtained individual cell annotations derived from the original study based on the acquired single-cell data. This approach ensured a comprehensive and accurate analysis of the cellular landscape across the dataset. Immune-related genes were obtained from the ImmPort (https://immport.niaid.nih.gov)^36^ and TimiGP R package^14^ (Supplementary Table 3).

### Clinical specimen collection and RNA sequencing

We assembled an in-house ICI cohort consisting of 14 patients diagnosed with metastatic or advanced NSCLC. These individuals underwent treatment with anti-PD-1/PD-L1 antibodies, either as monotherapy or in combination with anti-CTLA-4 antibodies, during the period from January 2018 to October 2022. Ethical approval for the collection of NSCLC tissue samples prior to ICI therapy was granted by the Medical Ethics Committee of Changhai Hospital. These samples are subsequently sent to Oncocare Inc., located in Suzhou, China, for RNA sequencing.

### Clinical outcomes

The primary endpoints for clinical evaluation were the Objective Response Rate (ORR) and Overall Survival (OS). The ORR was assessed in accordance with the Response Evaluation Criteria in Solid Tumors (RECIST) v1.1 guidelines^37^ for all cohorts, excluding the Hugo 2016 group^24^, where the assessment was performed using the immune-related RECIST (irRECIST) criteria. Patients were categorized into responder and non-responder groups: those attaining Complete Response (CR) or Partial Response (PR) were classified as responders, while individuals exhibiting Stable Disease (SD) or Progressive Disease (PD) were designated as non-responders.

### Multicolor Fluorescence Immunohistochemistry Protocol

We employed the Akoya OPAL Polaris 4-Color Manual IHC Kit (NEL811001KT) for multicolor immunofluorescence staining of tissue samples. The procedure commenced with baking the tissue slides to ensure sample adherence, followed by dewaxing to eliminate paraffin. Antigens were retrieved by heating the slides, and non-specific antibody binding was blocked. The slides were subsequently incubated with primary antibodies: CD4 (1:500, Cat# Ab133616, Abcam, USA), CD8A (1:2000, Cat# ab217344, Abcam, USA), and CD20 (1:100, Cat# ab64088, Abcam, USA). This was followed by incubation with HRP-conjugated secondary antibodies. For each antigen, signal amplification was achieved using tyramide signal amplification (TSA), and nuclei were stained with DAPI. Finally, coverslips were mounted for microscopy analysis, completing the staining process to facilitate detailed visualization of the cellular and molecular structures within the tissue samples.

### The framework of the AGAE score

The AGAE score was established to predict the clinical outcomes for cancer patients receiving ICI therapy. We firstly performed feature selection and engineering, including the gene-pairing process and ABESS algorithm, and then used the GA to find the best basic learner combination, which enabled us to construct the AGAE score, before this model was used to perform further analysis. For the construction and validation of the AGAE score and the exploration of the internal and external immune landscapes related to the AGAE score, please refer to the Supplementary Methods.

### CRISPR screening data

To investigate potential therapeutic targets of AGAE score, we aggregated data from seven published CRISPR/Cas9 screening studies that evaluated the individual effects of each gene knockout on tumor immunity, including Freeman^38^, Kearney^39^, Manguso^40^, Pan^41^, Patel^42^, Vredevoogd^43^, and Lawson^44^ (Supplementary Table 4). The first six CRISPR studies were previously curated by Fu et al.^45^. In addition to Fu et al., we further compiled another CRISPR cohort from Lawson et al.^44^. Based on model cell lines and treatment conditions, these seven CRISPR studies were categorized into 17 datasets. The CRISPR analysis encompasses melanoma, breast cancer, colon cancer, and renal cancer cell lines. We utilized these data to identify genes that are more likely to modulate lymphocyte-mediated cancer killing and influence immunotherapy responses across different datasets. The CRISPR screening process involves genome-wide CRISPR-Cas9 knockout in cancer cell lines under various conditions, followed by RNA-seq to estimate sgRNA abundance. Log-fold changes of sgRNA reads are calculated to assess gene knockout effects on cancer fitness under immune pressure. Normalized z-scores derived from these changes eliminate batch effects and enable cross-study comparisons. Lower z-scores indicate better immune responses post-knockout. Genes are ranked based on average z-scores across 17 datasets, with top-ranked genes (lower z-scores) characterized as immune resistant. For the identification and validation of key genes driven by the AGAE score, please refer to the Supplementary Methods.

### Statistical analysis

Pearson’s or Spearman’s coefficients were employed to assess correlations between variables. A t-test was utilized to analyze group differences when the variables followed a normal distribution; otherwise, the Mann-Whitney U test was applied. Differences among categorical variables were evaluated using Pearson’s chi-squared test or Fisher’s exact test. Kaplan-Meier (K-M) analysis was performed to generate survival curves, with the log-rank test used to assess significance. Receiver Operating Characteristic (ROC) curve analysis was conducted to establish optimal threshold values. Independent risk factors for clinical response were evaluated using multivariate logistic regression. A p-value of less than 0.05 was considered statistically significant. All statistical tests were two-sided, and all analyses were performed using R software (version 4.1.3).

## Results

### Development of AGAE score

Figure 1A illustrates the comprehensive workflow of the AGAE score development. Initially, feature processing was conducted on the training set. To mitigate batch effects, immune-related genes were paired, generating 187,6953 IRGPs. Multivariable logistic regression with cancer type adjustment (P < 0.001) and ROC curve analysis (AUC > 0.6) identified 2537 key IRGPs in the training cohort (Supplementary Table 5). Subsequently, to optimize predictive performance, four machine learning regression algorithms (ERN, RF, SVM, and KNN) were employed to construct basic learners in the training cohort. A 10-fold cross-validation and grid search were performed to account for parameter influence, ultimately yielding 50 basic learners (m1–m50) for further investigation (Supplementary Table 6). Utilizing a stacking strategy, the predicted values from basic learners in the training cohort served as input data to construct an ensemble model, termed the AGAE score. The ensemble model’s efficacy hinged on the diversity and accuracy of its basic learners. To automate this process, traditionally performed manually, we implemented a GA to optimize basic learner combinations, enhancing ensemble model accuracy. The final AGAE score incorporated ten models: m3, m5, m6, m16, m19, m23, m24, m29, m40, and m46. These comprised three ERN algorithm (m3, m5, and m6), two KNN algorithms (m40, m46), two RF algorithm (m16 and m19), and three SVM (m23, m24, and m29), demonstrating algorithmic diversity and validating the GA’s automation efficacy. The AGAE score was ultimately constructed in the training set using the predicted values from these four basic learners as input data, with stepwise logistic regression as the ensemble learning method.

**Figure 1.**
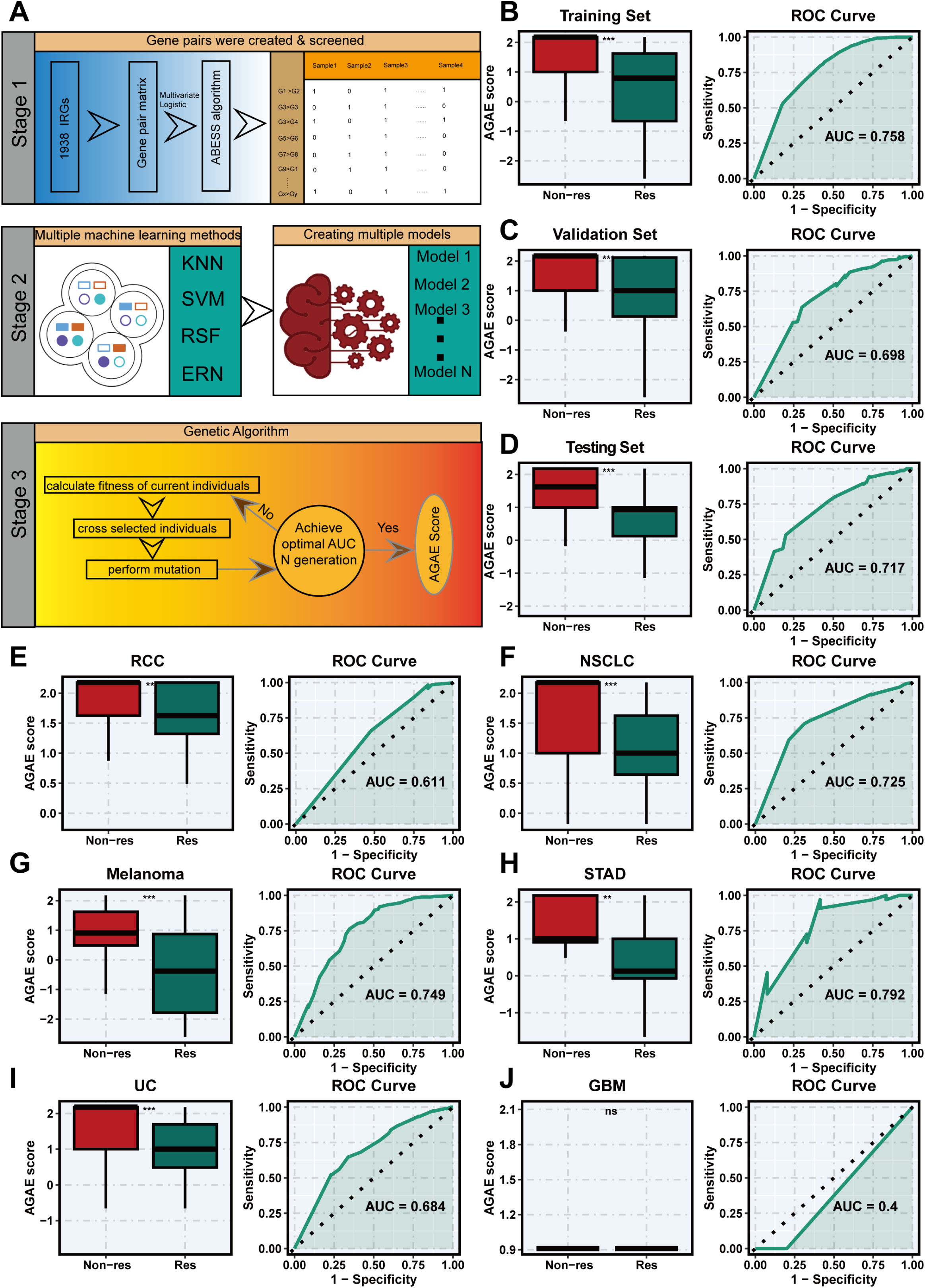
Prediction of ICI outcomes using AGAE score. **A.** Flow chart of training, validating, and testing the AGAE score constructed using genetic algorithms and ensemble learning. **B-D.** The distribution of AGAE score between responders and nonresponders of ICI therapy in the training (**B**), validating (**C**), and testing (**D**) cohorts (left). ROC curves of AGAE score to predict the benefits of ICI therapy in the training (**B**), validating (**C**), and testing cohorts (**D**) (right). **E-J.** The distribution of AGAE score between responders and nonresponders of ICI therapy in RCC (**E**), NSCLC (**F**), melanoma (**G**), STAD (**H**), UC (**I**), and GBM patients (**J**) (left). ROC curves of AGAE score to predict the benefits of ICI therapy in RCC (**E**), NSCLC (**F**), melanoma (**G**), STAD (**H**), UC (**I**), and GBM patients (**J**) (right).

### Assessment of AGAE score

To further evaluate the AGAE score’s efficacy in predicting clinical responses, we applied it to the training, validation, and test cohorts. The AGAE score demonstrated significant differentiation between response and non-response groups across all three cohorts. Notably, patients with lower AGAE scores exhibited a higher likelihood of benefiting from ICI therapy (all P < 0.001) (Figure 1B–D). ROC curve analysis was performed to assess the AGAE score’s predictive accuracy. The score achieved substantial AUC values in all cohorts: 0.758 for the training set, 0.698 for the validation set, and 0.717 for the test set (Figure 1B–D). Patients were stratified into high and low AGAE score groups using the optimal threshold from the training cohort. The high-AGAE score group showed significantly poorer OS across all three cohorts (P < 0.05) (Supplementary Figure 1A-C). Time-dependent ROC curves confirmed the AGAE score’s effectiveness in predicting OS for patients receiving ICI therapy (Supplementary Figure 1D-F). We further examined the AGAE score’s performance across various tumor types. In RCC, NSCLC, UC, melanoma, and STAD, the AGAE score was consistently lower in the response group (all P < 0.05), with ROC-AUC values exceeding 0.6 (Figure 1E–I). However, no significant difference was observed in GBM (Figure 1J), potentially due to the limited sample size for this tumor type. These findings underscore the AGAE score’s robust predictive capability across multiple cancer types, highlighting its potential as a valuable tool for guiding ICI therapy decisions.

### Comparison of AGAE score with Other Features and Webtool development

Recent advancements in sequencing technologies and machine learning algorithms have propelled researchers to develop predictive signatures for immunotherapy response. Numerous signatures, based on distinct biological characteristics and molecular types, have demonstrated predictive value for ICI therapy from various perspectives. We systematically evaluated 12 published signatures alongside the AGAE score. Comparative analysis revealed that the AGAE score exhibited superior predictive performance across the training, validation, and test cohorts (Figure 2A–C).

**Figure 2.**
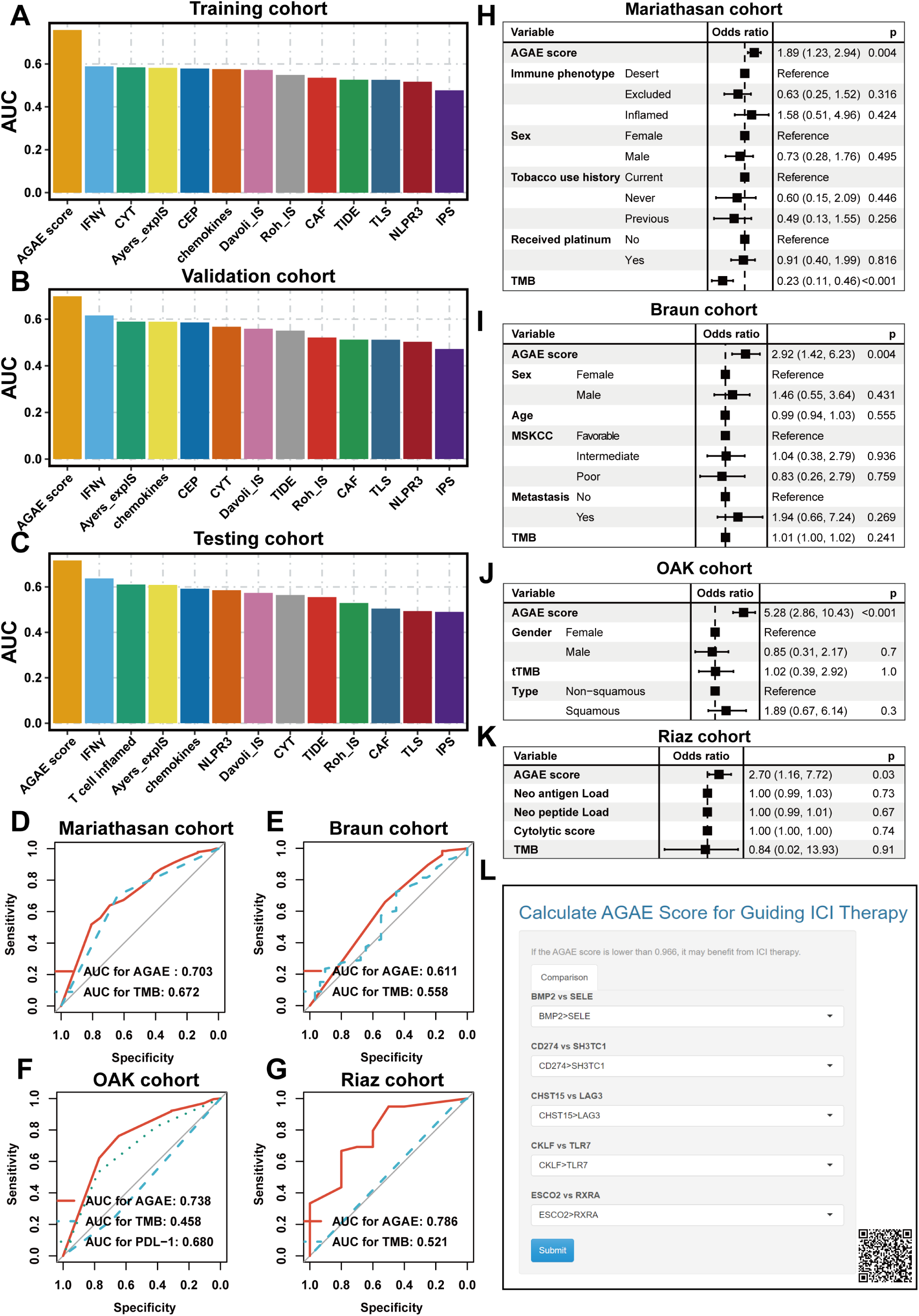
Assessment of AGAE score. **A-C.** Comparing AGAE scores with published signatures based on ROC Curves in the training (**A**), validating (**B**), and testing (**C**) cohorts. **D-G.** Comparing AGAE scores with TMB and PDL-1 based on ROC Curves in the Mariathasan (**D**), Braun (**E**), OAK (**F**),and Riaz (**G**) cohorts. **H-K.** Multivariate logistic regression analysis in the Mariathasan (**H**), Braun (**I**), OAK (**J**),and Riaz (**K**) cohorts. **L.** The page of AGAE score web-tool.

Leveraging the comprehensive clinical data available in the Mariathasan, Braun, OAK, and Riaz cohorts, we conducted ROC curve analyses to compare the AGAE score’s predictive performance against PD-L1 expression and TMB. The AGAE score consistently outperformed both PD-L1 and TMB in these cohorts (Figure 2D–G). Multivariate logistic regression analysis further confirmed the AGAE score as an independent predictor of clinical response (all P < 0.05) (Figure 2H–K). Recognizing the AGAE score’s robust predictive capability for ICI therapy response, we developed a user-friendly web tool to enhance its accessibility for the research community. This tool is illustrated in Figure 2L and can be accessed at https://ici-theaphy-gms.shinyapps.io/my_shiny_app/. These findings underscore the AGAE score’s potential as a valuable clinical decision-making tool for ICI therapy, offering improved predictive accuracy compared to existing biomarkers and signatures.

### Intrinsic Immune Landscapes Underlying the AGAE Score

To further elucidate the relationship between the immune landscape and AGAE score, we conducted multiomics analysis of TCGA cohort. Applying the formula and cutoff derived from the training cohort, we stratified the TCGA cohort into high AGAE score and low AGAE score groups (Figure 3A). Genomic analysis revealed significantly larger leukocyte, lymphocyte, and tumor-infiltrating lymphocyte (TIL) fractions in the low AGAE score group (P < 0.001) (Figure 3B-D). Corroborating these findings, TIL fraction estimates from H&E-stained slides using deep learning methods showed consistent results (P < 0.001) (Figure 3E). The low AGAE score group exhibited a significantly higher proportion of immune-stimulatory cells, particularly CD8^+^ T cells (P < 0.001) (Figure 3F). Further analysis using immune infiltration scores from Danaher et al. (Figure 3G) and immune signature scores (Figure 3H) confirmed the greater abundance of immune cells in the low AGAE score group (all P < 0.05). Unsupervised clustering of TCGA cohort patients based on immune signature scores revealed two distinct immune patterns: high and low immune infiltration (Figure 3I). Notably, the high immune infiltration group was significantly enriched with cases from the low AGAE score group (P < 0.001) (Figure 3J). These findings collectively demonstrate a strong association between the AGAE score and the immune microenvironment, with the low AGAE score group characterized by enhanced immune cell infiltration.

**Figure 3.**
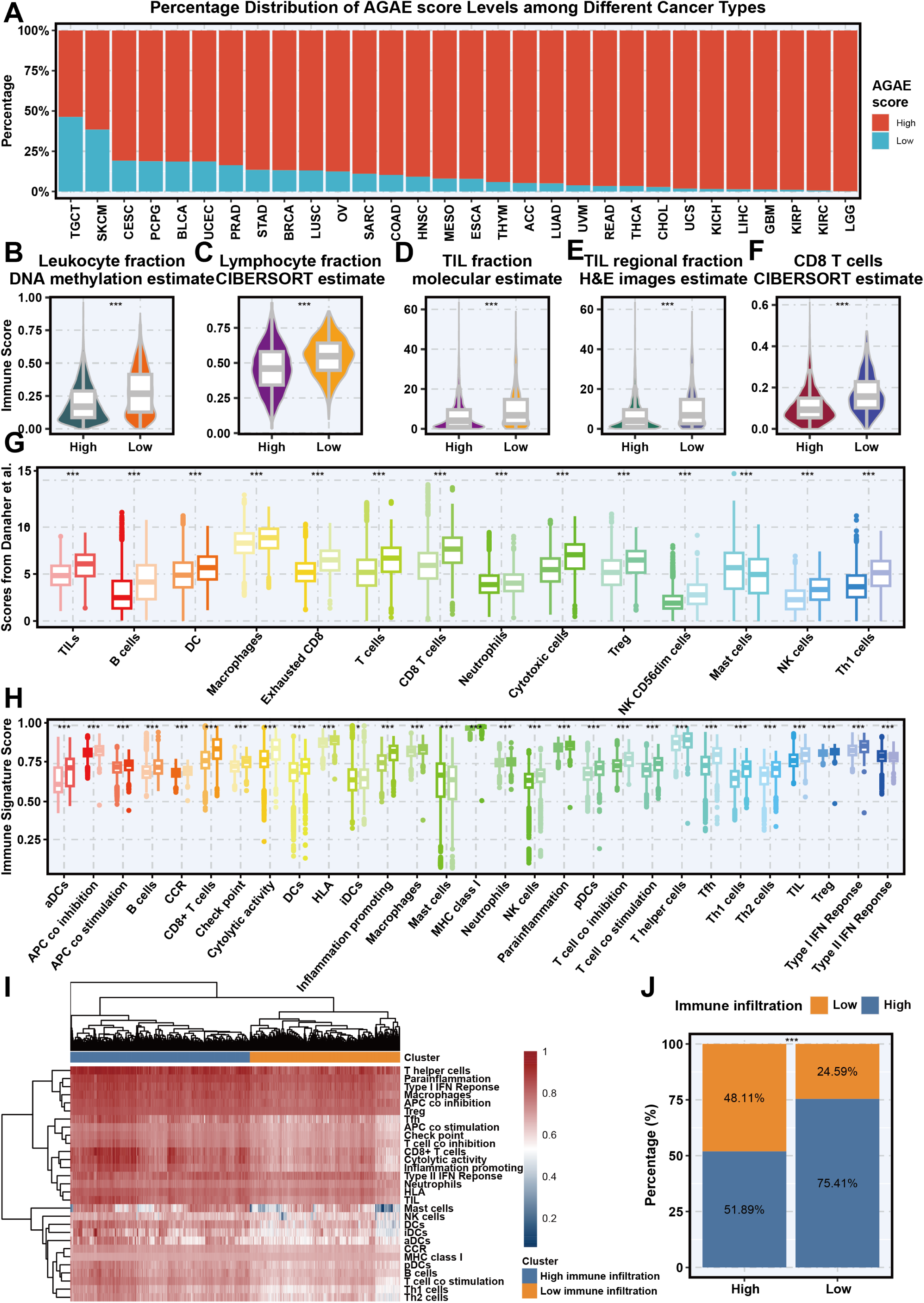
Immune landscapes of high AGAE score and low AGAE score groups in the TCGA Cohort. **A.** AGAE score for 30 cancers. **B.** Comparison of leukocyte fractions based on DNA methylation data between the high AGAE score and low AGAE score groups. **C.** Comparison of lymphocyte fractions estimated by the CIBERSORT method based on RNA-sequencing data between the high AGAE score and low AGAE score groups. **D.** Comparison of TIL fractions based on molecular estimates from the processing of cancer genomics data between the high AGAE score and low AGAE score groups. **E.** Comparison of TIL regional fractions based on estimates from processing diagnostic H&E images between the high AGAE score and low AGAE score groups. **F.** Comparison of CD8^+^ T cells estimated by the CIBERSORT method based on RNA-sequencing data between the high AGAE score and low AGAE score groups. **G.** Comparison of 14 immune cells estimated by the Danaher method based on RNA-sequencing data between the high AGAE score and low AGAE score groups. **H.** Comparison of the 29 immune signatures estimated by the ssGSEA method based on RNA-sequencing data between the high AGAE score and low AGAE score groups. **I.** Unsupervised clustering based on 29 immune signatures in the cohort from TCGA, yielding two stable immune subtypes. **J.** The proportions of high immune infltration and low immune infltration estimated by 29 immune signatures in the high AGAE score and low AGAE score group. *p < 0.05; ***p < 0.001.

In the low AGAE score group, immune signature scores were markedly higher at tumor sites compared to normal sites. Conversely, the high AGAE score group exhibited lower immune signature scores at tumor sites than at normal sites (Figure 4A). The low AGAE score group demonstrated significantly stronger correlations among immune activities compared to the high AGAE score group (Figure 4B, C). Gene Set Enrichment Analysis (GSEA) identified 33 significantly enriched pathways in the low AGAE score group, including 19 immune-related pathways such as "natural killer cell mediated cytotoxicity" (all p -adjust < 0.05) (Figure 4D). The high AGAE score group showed no enrichment in immune-related pathways. Low AGAE score tumors were associated with significantly higher CYT scores (P < 0.001) (Figure 4E), while the high AGAE score group exhibited a significantly larger abundance of fibroblasts (P < 0.05) (Figure 4F). These findings suggest that the low AGAE score group is characterized by abundant immune cells at tumor sites, potentially contributing to improved response to immune checkpoint inhibitor ICI therapy. Conversely, the prevalence of fibroblasts in the high AGAE score group may contribute to extrinsic immune escape mechanisms. The low AGAE score group showed higher chemokine expression (most P < 0.05) (Figure 4G), aligning with increased immune cell infiltration (Figure 3). This suggests chemokine enrichment may drive the enhanced immune response in the low AGAE scoregroup.

**Figure 4.**
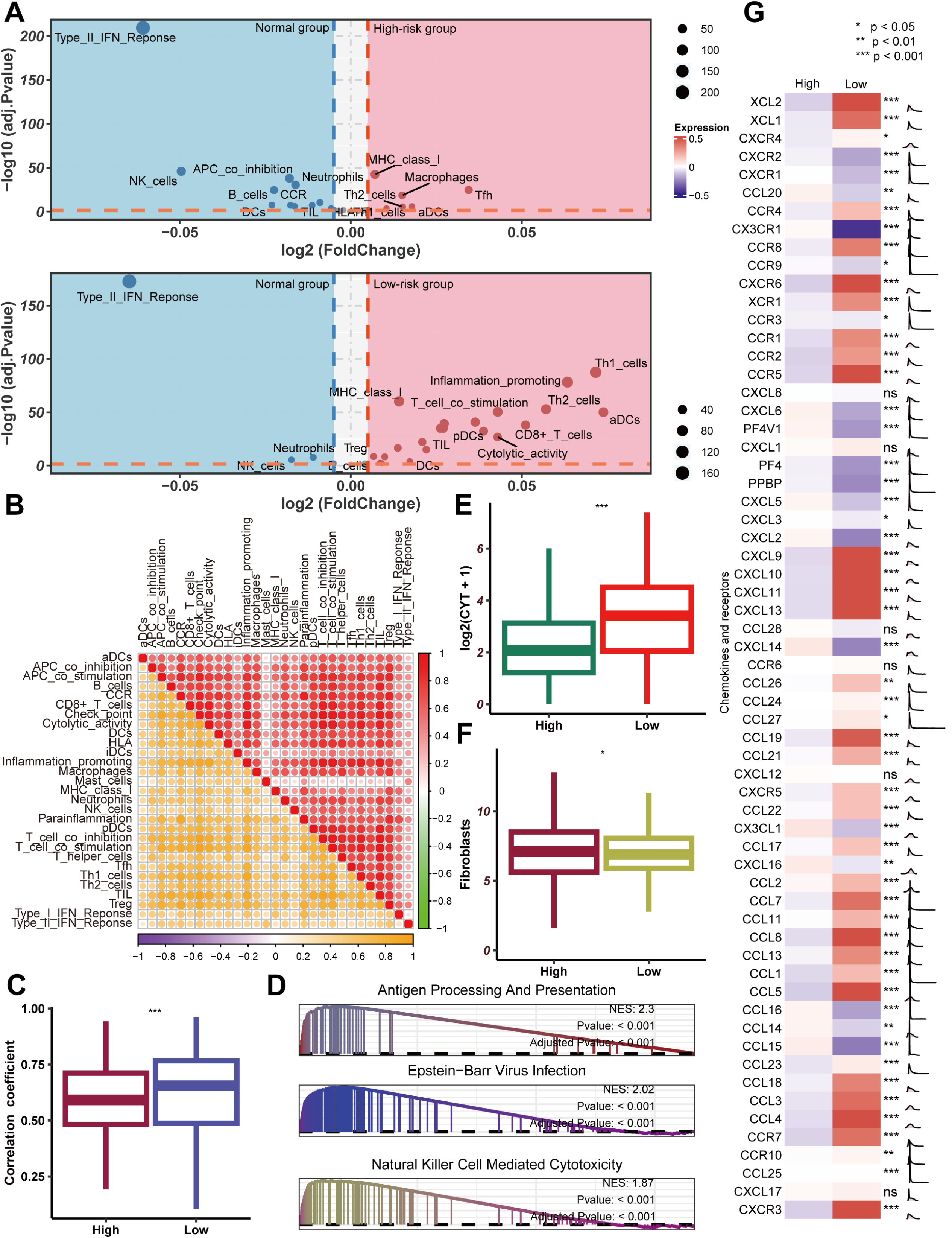
Potential extrinsic immune response landscapes in the high AGAE score and low AGAE score groups. **A.** Volcano plots of 29 immune signatures in the high AGAE score and low AGAE score groups. Immune signatures enriched in cancer tissues are marked in red; immune signatures enriched in normal tissues are marked in blue. **B.** Correlations among 29 immune signatures in the high AGAE score (top right panel) and low AGAE score (low left panel) groups. **C.** Comparison of the correlation coefcient among 29 immune signatures between the high AGAE score and low AGAE score groups. **D.** GSEA of the high AGAE score and low AGAE score groups. **E.** Comparison of the CYT score between the high AGAE score and low AGAE score groups. **F.** Comparison of fbroblast abundances between the high AGAE score and low AGAE score groups. **G.** Comparison of the expression patterns of chemokines between the high AGAE score and low AGAE score groups.

### Intrinsic Immune Landscapes Underlying the AGAE Score

We initially examined key determinants of tumor immunogenicity between the two groups. The low AGAE score group exhibited elevated mutation rates and neoantigen burdens compared to the high AGAE score group (all P < 0.05) (Figure 5A). Additionally, significantly higher TCR and BCR diversity was observed in the low AGAE score group (all P < 0.001) (Figure 5A). The low AGAE score group demonstrated increased CNV burden and aneuploidy (all P < 0.001) (Figure 5A). The high AGAE score patients displayed lower intertumoral heterogeneity than their low AGAE score patients (P < 0.001) (Figure 5A). These features collectively suggest a more immunogenic tumor microenvironment, which is often predictive of better responses to ICIs therapy. The increased intertumoral heterogeneity in this group may also contribute to a broader range of potential neoantigens, potentially making these tumors more susceptible to immune recognition and attack. Therefore, patients in the low AGAE score group may be more likely to benefit from immunotherapy strategies.

**Figure 5.**
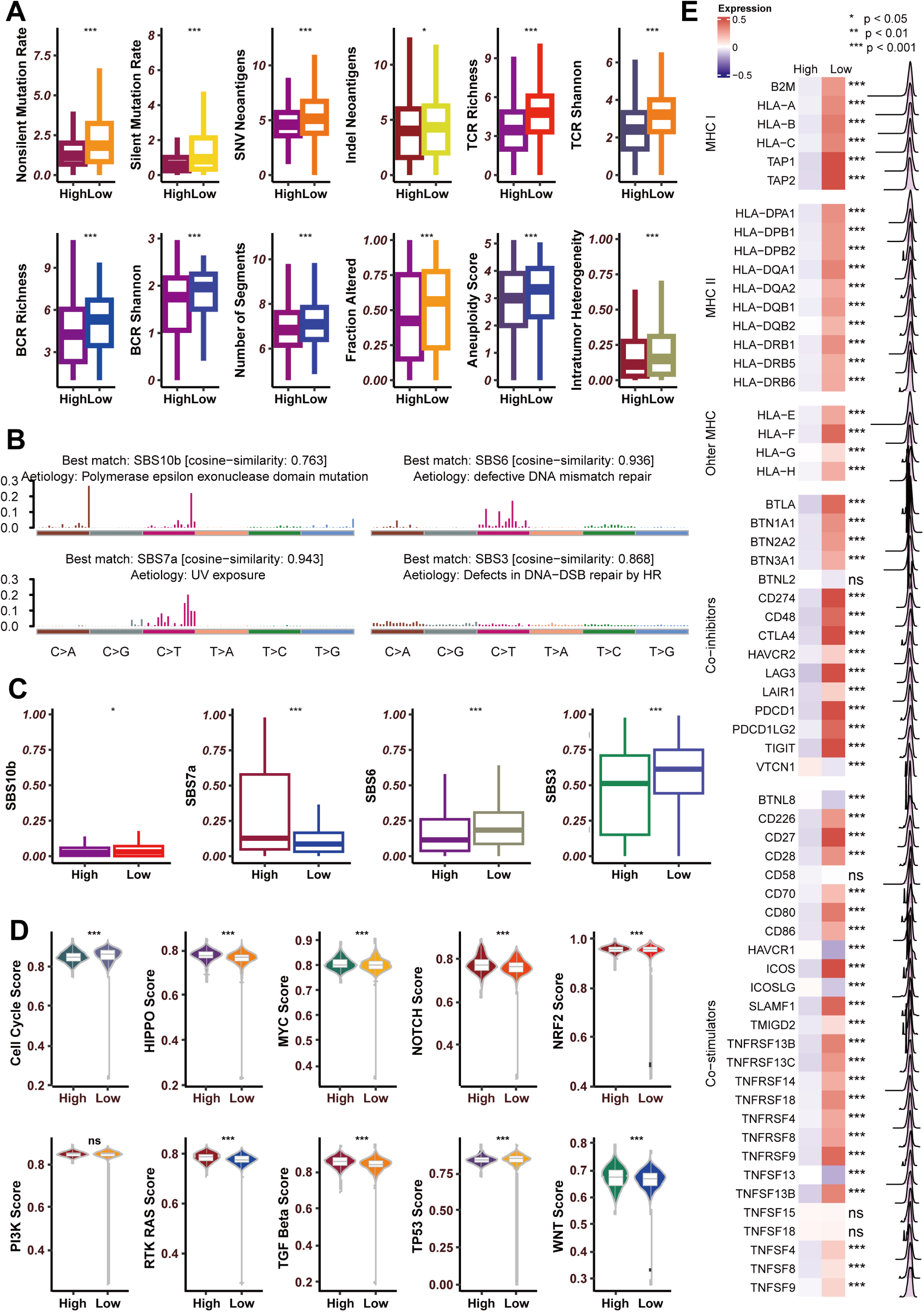
Potential intrinsic immune response and escape landscapes in the high AGAE score and low AGAE score groups. **A.** Comparison of immunogenomic indicators between the high AGAE score and low AGAE score groups. **B.** Mutational activities of four corresponding extracted mutational signatures. **C.** Comparison of four mutational signatures between the high AGAE score and low AGAE score groups. **D.** Comparison of enrichment scores of 10 oncogenic pathways between the high AGAE score and low AGAE score groups. **E.** Comparison of the expression patterns of MHC molecules, costimulators and coinhibitors between the high AGAE score and low AGAE score groups.

To elucidate the mutational landscape in both groups, we conducted an in-depth analysis of somatic mutation data from the TCGA dataset. This investigation revealed four distinct mutational signatures: SBS10b, SBS7a, SBS6, and SBS3 (Figure 5B). Notably, SBS10b, SBS3, and SBS6 exhibited significantly higher frequencies in the low AGAE score group compared to their high AGAE score counterparts (all P < 0.05) (Figure 5C). Of particular interest is the SBS6 signature, which is linked to deficiencies in DNA mismatch repair (MMR) mechanisms and has been implicated in modulating immune responses^46^.

We subsequently calculated enrichment scores for oncogenes in 10 common oncogenic pathways across both groups. The cell cycle and TP53 pathways showed higher scores in the low AGAE score group, while the HIPPO, MYC, NOTCH, NRF2, RAS, TGF Beta and WNT pathways were enriched in the high AGAE score group (all P < 0.001) (Figure 5D). The WNT pathway has been implicated in immune exclusion^47^.

The high AGAE score group demonstrated reduced expression of MHC I- and II-related antigen-presenting molecules compared to the low AGAE score group (all P <0.001), suggesting intrinsic immune escape (Figure 5E). Conversely, the low AGAE score group exhibited elevated expression of most MHC genes (Figure 5E), indicative of enhanced immunogenicity. Furthermore, immune checkpoint molecules (e.g., PD-1, PD-L1, and CTLA4) and costimulatory molecules were more highly expressed in the low AGAE score group (most P<0.001) (Figure 5E). These findings suggest that these immune checkpoint molecules may contribute to the response to ICI therapy in the low AGAE score group.

### Copy number features of the AGAE score

Substantial chromosomal aberration disparities were observed between the high AGAE score and low AGAE score groups (Figure 6A). The low AGAE score group exhibited focal amplification peaks in well-characterized immune genes, including PD-L1 and PD-L2 (both at 9p24.1), contrasting with the high AGAE score group (Figures 6B-C). Venn diagram analysis revealed 319 shared genes in amplified chromosome regions across both groups, with 217 and 1225 genes uniquely amplified in the high AGAE score and low AGAE score groups, respectively (Figure 6D). Gene Ontology annotation of these uniquely amplified genes and subsequent clustering of the top 10 biological processes (Figure 6E) demonstrated significant enrichment of two immune-related processes in the low AGAE score group: "T cell costimulation" and "B cell proliferation". Conversely, the high AGAE score group showed significant enrichment in "regulation of fibroblast proliferation" but lacked enrichment in immune-related processes. This finding aligns with previous observations of increased immune cell presence in the low AGAE score group (Figures 3B-J) and elevated fibroblast levels in the high AGAE score group (Figure 4F). Notably, PD-L1 and PD-L2, located within the low AGAE score group-specific amplification peak at 9p24.1, were annotated in both immune-related biological processes, suggesting their potential significance in modulating immune status within this group (Figure 6F). Analysis of mRNA expression data from the TCGA cohort revealed significantly higher expression levels of PD-L1 and PD-L2 in the low AGAE score group (Figure 6G), corroborating the CNV data. These findings indicate that tumor-specific CNVs contribute to the observed differences in immune infiltration patterns.

**Figure 6.**
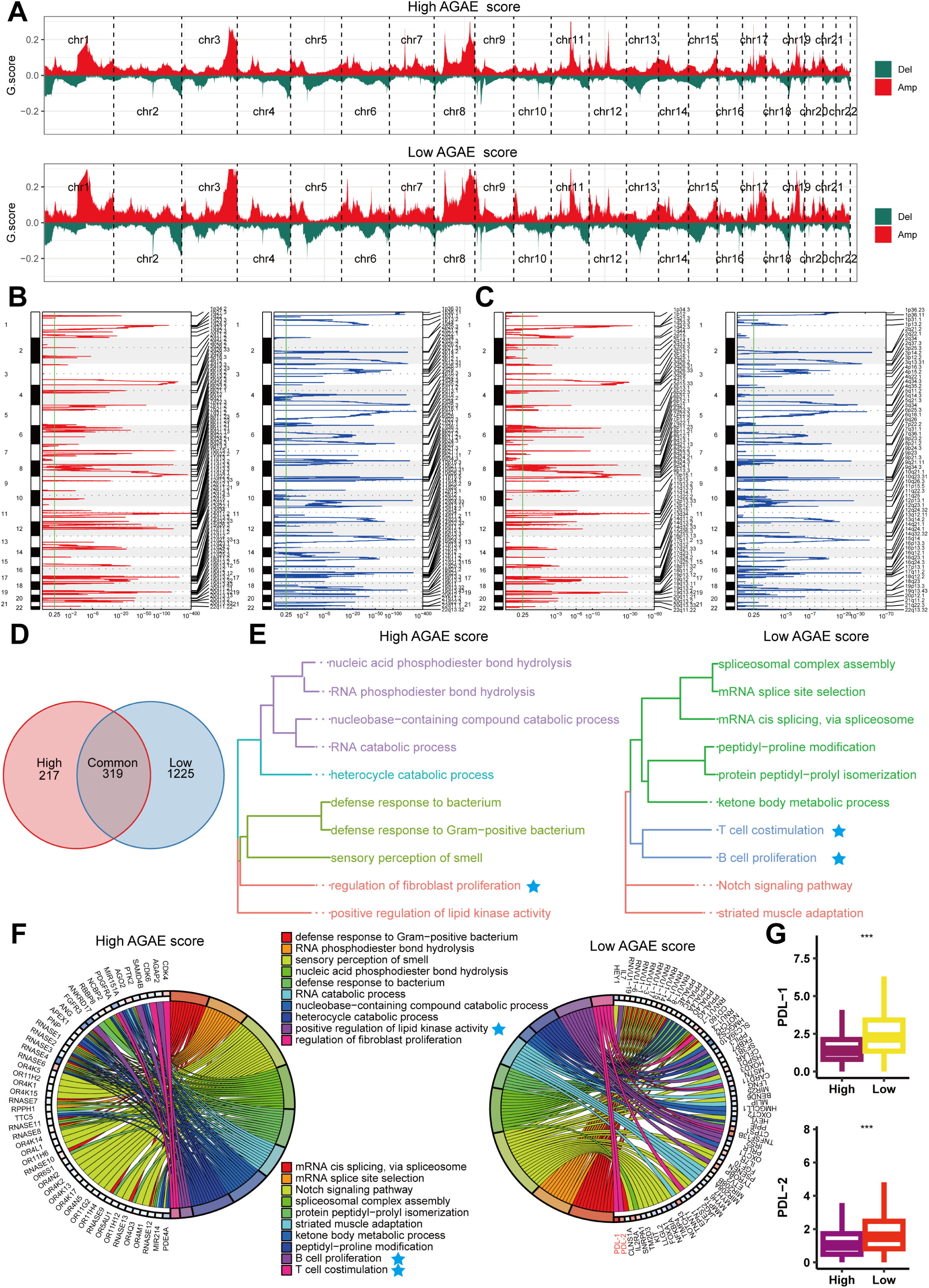
Copy number alterations in the high AGAE score and low AGAE score groups. **A.** Copy number profles of the high AGAE score (above) and low AGAE score (below) groups, with gains shown in red and losses shown in green. **B.** Detailed cytobands with focal amplifcation (left) and deletion (right) peaks identifed in the high AGAE score group. **C.** Detailed cytobands with focal amplifcation (left) and deletion (right) peaks identifed in the low AGAE score group. **D.** Venn diagrams showing signifcantly amplifed genes in the high AGAE score and low AGAE score groups. Each circle in the Venn diagram represents one group, and the number in the overlaid area represents common genes between the groups. **E.** Cluster analysis of the top 10 biological processes in the high AGAE score (left) and low AGAE score (right) groups. **F.** Circular plot of the 10 biological processes and corresponding enriched genes in the high AGAE score (left) and low AGAE score (right) groups. **G.** Comparison of the mRNA expression of PD-L1 and PD-L1 between the high AGAE score and low AGAE score groups in the cohort from TCGA.

### Validation of predictive value of the AGAE score in the in-house cohort

We performed multiplex immunofluorescence staining on NSCLC patient tissue samples using DAPI, CD4, CD8, and CD20 markers. Subsequent analysis demonstrated that specimens from the low AGAE score cohort exhibited significantly higher infiltration of CD4^+^ T cells, CD8^+^ T cells, and B lymphocytes compared to those from the high AGAE score group (Figure 7A). Post-immunotherapy imaging was conducted to evaluate patient responses according to RECIST criteria (Figures 7B). The predictive value of the AGAE score for neoadjuvant immunotherapy outcomes was confirmed through ROC analysis, yielding a robust AUC of 0.733 (Figure 7C and D). Additionally, the responder group included a substantial number of cases from the low AGAE score group (Figure 7E), although this did not reach statistical significance (P = 0.09), likely due to the limited sample size (n = 14).

**Figure 7.**
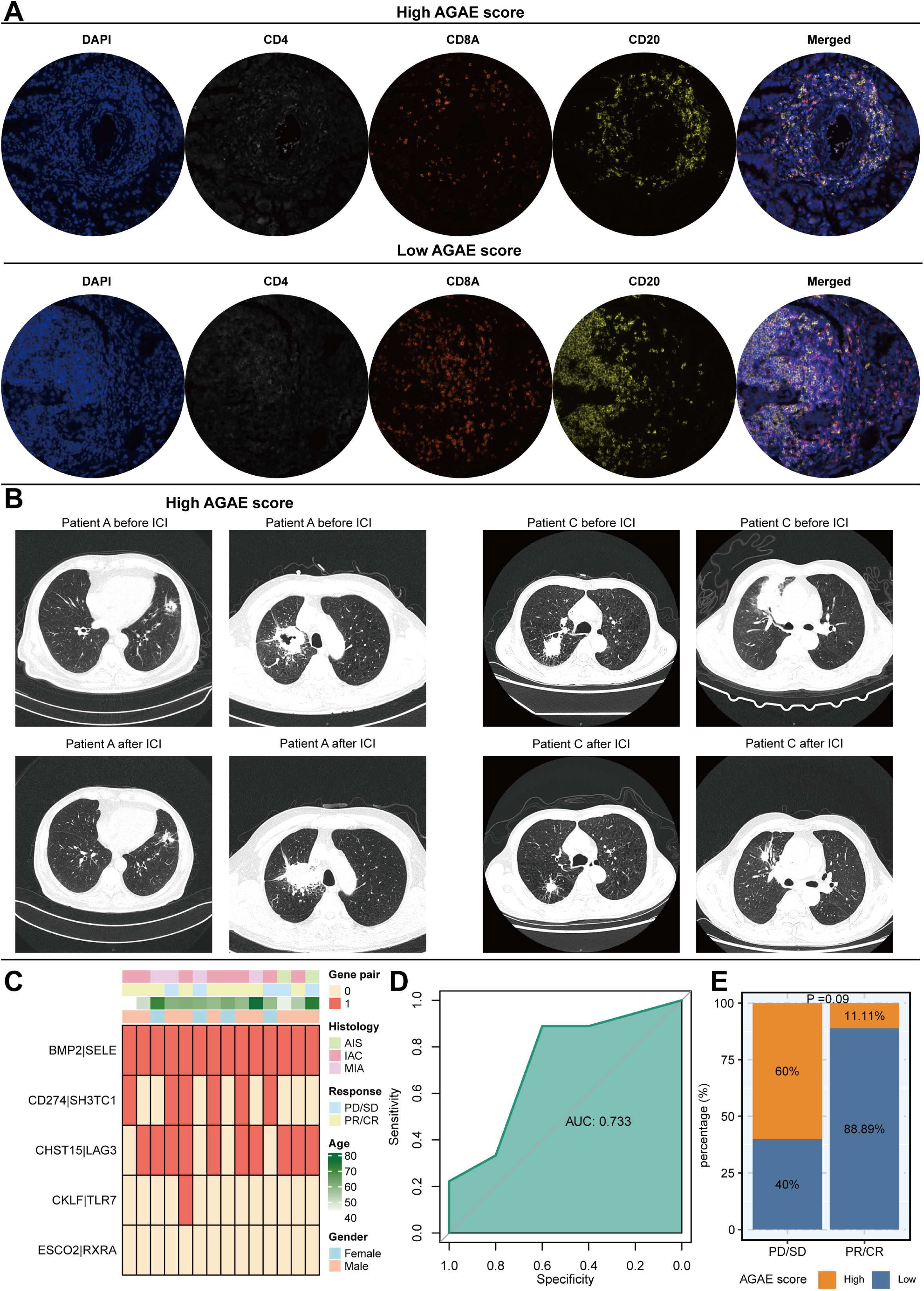
Predictive value of AGAE score in the in-house cohort. **A.** Representative images of DAPI, CD4, CD8, and CD20 based on multiplex IF staining in the high AGAE score group and low AGAE score group. **B.** Imaging manifestations before and after immunotherapy in the high-risk group and low AGAE score groups. **C.** Basic clinical characteristics of the in-house cohort. **D.** The ROC of AGAE score predicting clinical response in the in-house cohort. **E.** Differential analysis of the AGAE score between response and non-response groups.

### Exploration of potential therapeutic targets from AGAE score using CRISPR screen data

We conducted a comprehensive analysis of immune response data from seven CRISPR cohorts, encompassing 22,505 genes. These cohorts were further stratified into 17 distinct datasets based on model cell types and treatment conditions. Genes were ranked according to their mean z scores, with top-ranked genes identified as immune-resistant and bottom-ranked genes as immune-sensitive. Knockout of immune-resistant genes potentially enhances anti-tumor immunity, while disruption of immune-sensitive genes may attenuate it. Figure 8A illustrates the gene ranking methodology employed in this study. Then, we utilized univariate Cox regression on the meta cohort, which includes the integrated training set, validation set, and test set, to explore prognostic genes with a high AGAE score (P < 0.05). We then ranked these prognostic genes based on the CRISPR screen data, with CEP55 coming in first place (Figure 8B). As shown in Figure 8C, in TCGA pan-cancer datasets, the expression of CEP55 was upregulated in tumor samples compared to normal samples (most P < 0.05). Our investigation into CEP55’s potential biological functions utilized both TCGA pan-cancer datasets and scRNA-seq data. Analysis revealed CEP55’s predominant enrichment in cell cycle-related pathways, notably the G2M CHECKPOINT and E2F TARGETS (Figure 8D). Focusing on the pan-cancer scRNA-seq dataset, we isolated tumor epithelial cells for dimensionality reduction and clustering (Figure 8E). This approach demonstrated CEP55 expression in a subset of tumor epithelial cells (Figure 8F). GSEA further corroborated that CEP55 was also enriched in cell cycle-related pathways (Figure 8G). In conclusion, our findings position CEP55 as a promising target for enhancing the efficacy of ICI therapy. By simultaneously addressing tumor cell proliferation and immune evasion mechanisms, CEP55-targeted approaches could potentially overcome current limitations in immunotherapy and improve outcomes for cancer patients.

**Figure 8.**
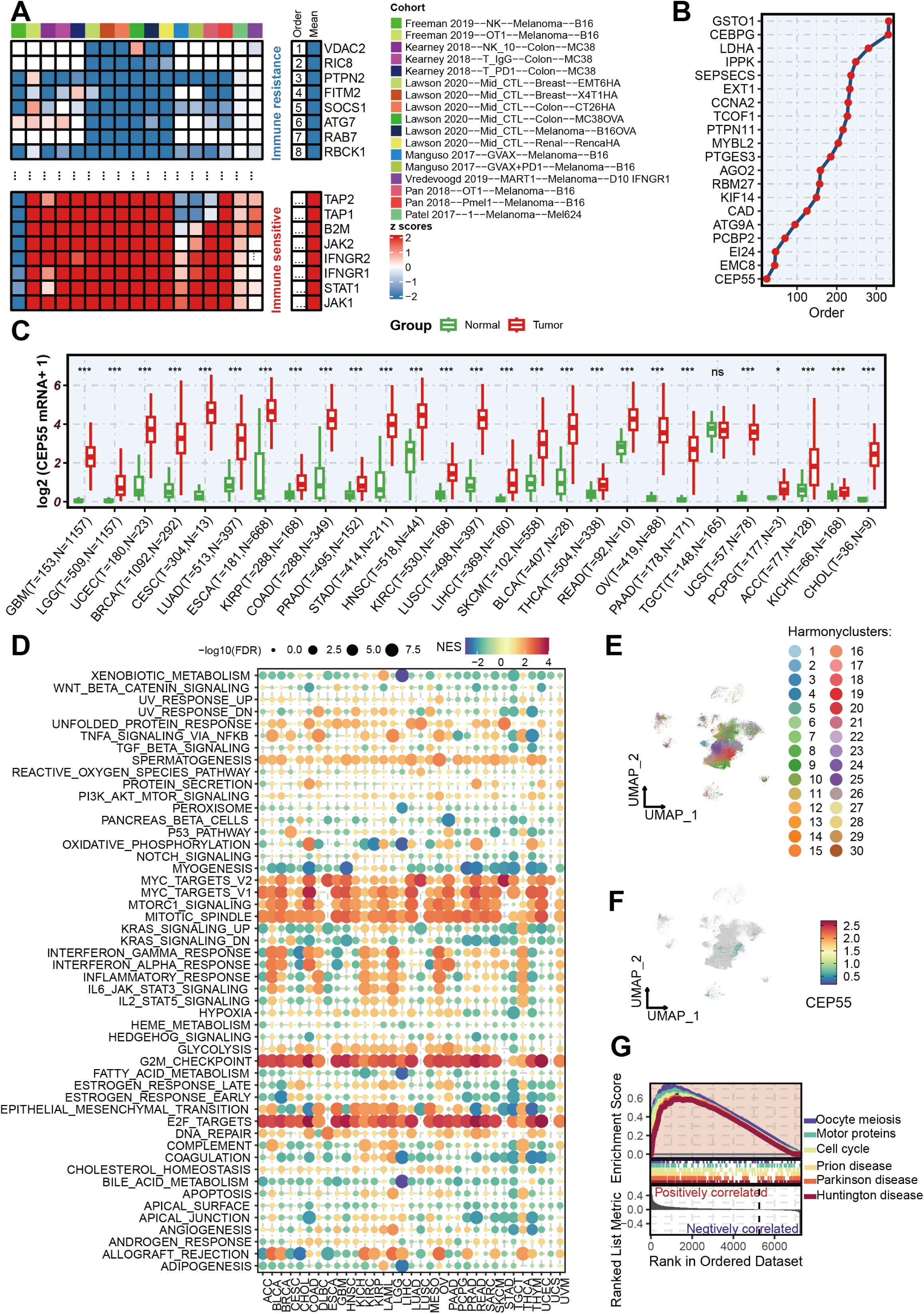
Exploration of potential treatment targets from AGAE score using CRISPR screening data. **A.** Ranking of genes based on their knockout efects on anti-tumor immunity across 17 CRISPR datasets. **B.** Univariate Cox regression analysis identified high AGAE score prognostic genes, with CEP55 ranking first based on CRISPR screen data. **C.** The expression of CEP55 across diverse cancers in the TCGA cohort. **D.** GSEA of CEP55 based on bulk RNA-seq datasets. **E-F.** The visualization of epithelial cells (**E**) and CEP55 mRNA expression (**F**) using Uniform Manifold Approximation and Projection (UMAP). **G.** GSEA of CEP55 based on scRNA-seq datasets. ^NS,^ no significant; *p < 0.05; **p < 0.01; ***p < 0.001.

### Knockdown of CEP55 Suppresses Cancer Cell Malignant Behavior

To investigate the role of CEP55 in the malignant biological behavior of cancer cells, Tumor cells were transfected with control shRNA (shNC) or shRNA targeting CEP55 (shCEP55-1, shCEP55-2) (Supplementary Figure 2A). Colony formation assays showed that knockdown of CEP55 significantly reduced the ability of tumor cells to form colonies (P < 0.05) (Figures 9A and B). Furthermore, transwell migration and invasion assays indicated that CEP55 knockdown significantly inhibited the migration and invasion abilities of CT26 and 3LL cells (all P < 0.05) (Figures 9C-F). Flow cytometry results demonstrated that the proportion of apoptotic cells in CT26 and 3LL cells significantly increased after CEP55 knockdown (all P < 0.05) (Figures 9G and H). These findings suggest that CEP55 knockdown significantly inhibits colony formation, migration, and invasion of cancer cells, while promoting apoptosis. In summary, CEP55 knockdown significantly inhibits the malignant biological behavior of tumor cells.

**Figure 9.**
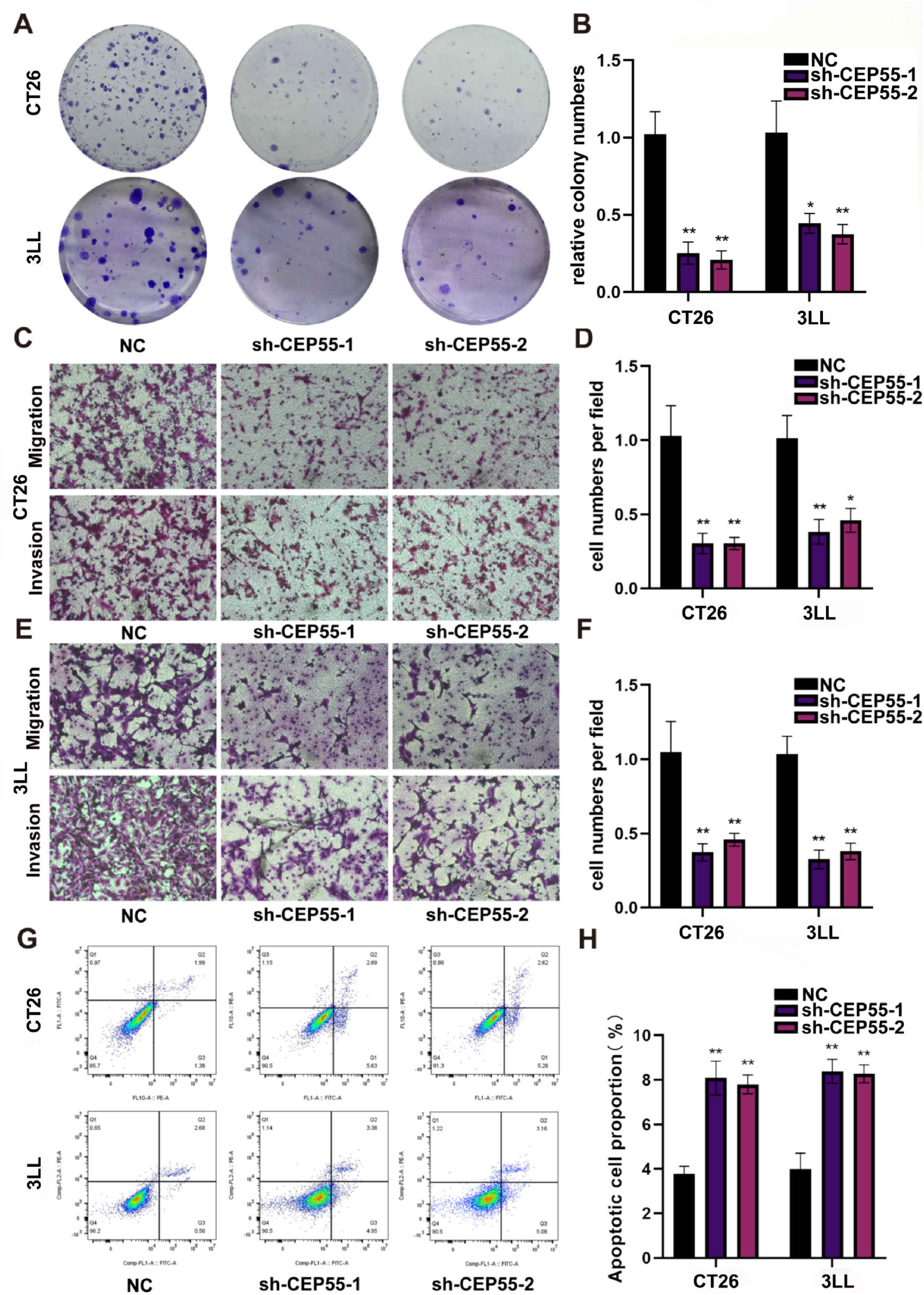
Knockdown of CEP55 suppresses cancer cell malignant behavior. **A-B.** Colony formation assays demonstrating that CEP55 knockdown notably inhibits cancer cell proliferation. **C-F.** Transwell assays evaluating the migration and invasion capabilities of CEP55-knockdown CT26 and 3LL cells. **G-H.** Flow cytometry evaluating that the proportion of apoptotic cells in CT26 and 3LL cells after CEP55 knockdown.

### CEP55 Downregulation Reversing CD8^+^ T Cell Exhaustion and Boosting ICT Efficacy in Mouse Subcutaneous Model

This study examined the effect of CEP55 knockdown on CD8^+^ T cell immune function in co-culture systems with CT26 and 3LL cancer cells. Tumor cells were transfected with either a non-targeting control shRNA (shNC) or CEP55-targeting shRNAs (shCEP55-1, shCEP55-2) and co-cultured with CD8^+^ T cells. Results demonstrated a significant increase in the percentage of IFN-γ^+^ CD8^+^ T cells in the CEP55 knockdown groups compared to the control group (P < 0.05) (Figures 10A and B). Similarly, the percentage of TNF-α^+^ CD8^+^ T cells was significantly elevated in both CT26 and 3LL cells following CEP55 knockdown (P < 0.05) (Figures 10C and D). Flow cytometry further revealed that CEP55 knockdown in tumor cells reduced the expression of exhaustion markers PD-1 and TIM-3 on CD8^+^ T cells, suggesting that decreased CEP55 expression reverses CD8^+^ T cell exhaustion and restores their function (P < 0.05) (Figures 10E and F). To assess the impact of CEP55 knockdown on tumor growth and survival, we utilized a mouse subcutaneous tumor model with 3LL cells (n = 5 per group). In this model, CEP55 knockdown (shCEP55-1) led to a reduction in tumor weight and volume compared to the control (shNC) group (Figures 10G, 10H, 10J). Kaplan-Meier survival analysis showed that CEP55 knockdown, particularly when combined with anti-PD1 therapy, significantly improved survival rates in 3LL tumor-bearing mice (P < 0.05) (Figure 10I). These findings suggest that CEP55 is a potential therapeutic target for enhancing immunotherapy efficacy.

**Figure 10.**
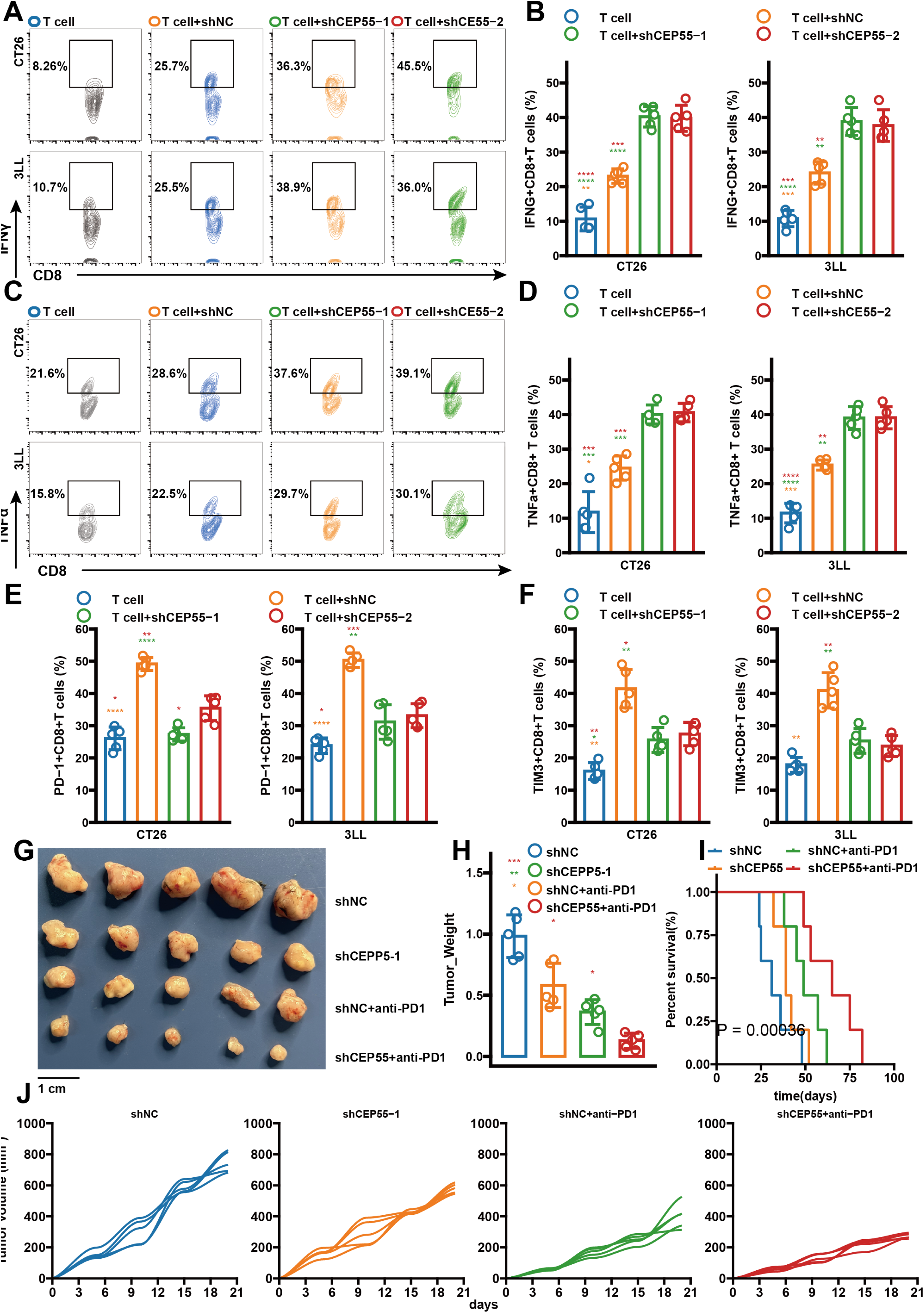
CEP55 knockdown enhances CD8^+^ T cell function and inhibits tumor growth. **A.** Flow cytometry analysis of IFN-γ^+^ CD8^+^ T cells co-cultured with CT26 or 3LL tumor cells transfected with shNC or shCEP55 (shCEP55-1, shCEP55-2). **B.** Quantification of IFN-γ^+^ CD8^+^ T cells in CT26 and 3LL co-culture systems. **C.** Flow cytometry analysis of TNF-α^+^ CD8^+^ T cells in the same co-culture systems. **D.** Quantification of TNF-α^+^ CD8^+^ T cells in CT26 and 3LL co-culture systems. **E-F.** Expression levels of exhaustion markers PD-1 and TIM-3 on CD8^+^ T cells after co-culture with CEP55 knockdown tumor cells. **G.** Representative images of tumors from the 3LL subcutaneous mouse tumor model with shNC, shCEP55, and shCEP55 combined with anti-PD1 treatments. **H.** Tumor weight in the 3LL tumor model across different treatment groups. **I.** K-M survival curves of CT26 tumor-bearing mice receiving shNC, shCEP55, and anti-PD1 treatments. **J.** Tumor growth curves (volume in mm³) for 3LL tumors in the respective treatment groups.

## Discussion

Robust predictive biomarkers are crucial for guiding immunotherapy decisions, optimizing resource allocation, and expediting clinical trials and regulatory approvals. While several biomarkers have been explored and implemented to forecast treatment outcomes, their predictive efficacy remains suboptimal^6–9^. This limitation underscores the urgent need for novel responsiveness indicators. Our study, encompassing 1,242 patients with diverse malignancies, introduces and validates the AGAE score—a five gene-pairs model designed to prognosticate clinical outcomes in patients receiving ICI therapy. To our knowledge, this represents the first pan-cancer immunotherapy predictive model leveraging ensemble learning that integrates both gene-pair and GA approaches.

Our study offers several innovative aspects and practical applications. First, our investigation encompassed diverse tumor types, including NSCLC, melanoma, and RCC. These malignancies represent the most prevalent cancer types currently treated with ICI therapy in clinical settings. Second, numerous mRNA-based signatures have emerged as potential predictors of immunotherapy efficacy. However, implementing multi-biomarker predictive models necessitates a comprehensive understanding of factors influencing the precision and accuracy of high-throughput assays in clinical settings. Paramount among these are sources of measurement variability, categorized into preanalytical (sample-intrinsic) and technical (platform-intrinsic) factors. Tissue-specific variations in mRNA expression are typically mitigated through the incorporation of multiple reference genes, with relative quantification employed to normalize target gene expression. Notably, the risk-score algorithms and cutoff values established for these mRNA signatures are often incompatible with alternative measurement methodologies, limiting their broad applicability across diverse data types^48^. To overcome this challenge, we adopted the gene-pairing method, specifically using IRGPs, which effectively eliminates batch effects. This approach not only benefits further ensemble learning models but also offers several key advantages. IRGPs help mitigate the "prognostic bias" caused by immune cell co-infiltration and capture the relative abundance between immune cells, potentially offering better insights into cancer behaviors. Gene pair-based predictive models demonstrate improved robustness and accuracy across different sequencing platforms and sample types, enhancing their translational potential. They also enable the detection of subtle differences between immune cell types and are less affected by technical variations in gene expression measurements. By addressing the batch effect issue and leveraging these advantages, this approach offers improved robustness, biological interpretability, and clinical applicability compared to using individual gene expression levels, particularly valuable for studying complex systems like the tumor immune microenvironment^14^. Third, our approach leverages genetic algorithms to optimize the selection of base learners within the ensemble model, thereby augmenting predictive accuracy. This integrated ensemble learning methodology demonstrates versatility beyond clinical efficacy prediction, extending to various analytical tasks where gene expression and clinical data are available. For instance, it holds promise in prognostic forecasting across diverse cancer types. Fourth, the AGEA score offers practical advantages in clinical application. It potentially shields non-responsive patients from immune-related adverse effects while facilitating prompt allocation to more suitable treatment options. Given that a typical treatment course averages $120,000^49^, implementing biomarker strategies that enhance diagnostic precision could yield substantial cost savings for interventions with anticipated diminished efficacy.

We utilized the multifaceted TCGA dataset to elucidate immunotherapy response patterns across cancer types. Our analysis revealed that the low AGAE score group exhibited an inflammatory immune profile, characterized by elevated CD8^+^ T cell infiltration and immunogenicity. Employing immune infiltration scores from Danaher et al. and ssGSEA methodologies, we observed significantly higher overall immune cell infiltration in the low AGAE score group, corroborating its robust antitumor immune activity. Numerous studies have established a positive correlation between TIL density and immune responses across various malignancies^50^. The low AGAE score group not only demonstrated increased cytotoxic T cell infiltration but also exhibited overexpression of immune checkpoint molecules, including PD-L1, PD-1, and CTLA-4, relative to the high AGAE score cohort. This combination of activated antitumor immunity, elevated checkpoint molecule expression, and heightened tumor immunogenicity likely underpins the enhanced responsiveness to ICI therapy observed in the low AGAE score group.

Given the strong performance of the AGAE score in predicting immunotherapy outcomes, there is an urgent need to identify potential targets associated with it. To this end, we utilized CRISPR datasets and ranked genes based on their log fold changes (logFCs) of sgRNA reads under immune-competent and immune-deficient conditions. Then, we utilized univariate Cox regression on the meta cohort to explore prognostic genes with a high AGAE score. This analysis identified CEP55 as a key gene for further investigation. CEP55, a member of the centrosome protein family and an expression product of immune escape-related genes, plays a crucial role in regulating mitosis and cytoplasmic division^51 52^. Its primary function involves anchoring microtubules, polymerizing associated proteins, and participating in spindle formation, which collectively regulate cell proliferation. Studies have indicated that CEP55 is overexpressed in various tumors^52 53^. The overexpression of CEP55 directly interacts with the p110 catalytic subunit of PI3K, leading to the upregulation of the PI3K/AKT pathway, which promotes cell transformation, proliferation, epithelial-to-mesenchymal transition, invasion, and migration, thereby facilitating cancer progression^54 55^. Notably, a previous functional genomic screening identified CEP55 as one of 182 genes whose perturbation could alter tumor sensitivity to T cell-mediated killing therapies^44^. In colorectal cancer, CEP55 overexpression is associated with immune exclusion and tumor progression, while its inhibition has been shown to enhance anti-tumor immunity and sensitivity to ICIs^56^. Our findings further revealed that CEP55 knockdown significantly inhibited the malignant biological behaviors of cancer cells and improved the efficacy of immunotherapy. Our study extends these findings by demonstrating that CEP55 knockdown not only suppresses the malignant potential of cancer cells but also potentiates the effectiveness of immunotherapies, highlighting CEP55 as a promising therapeutic target for cancer treatment.

### Limitations

Our investigation acknowledges several constraints that warrant consideration. Primarily, the 11 RNA-Seq cohorts utilized in our analysis encompass a limited spectrum of six cancer types. While the consistent inverse relationship between AGAE score and anti-tumor immunity, observed across a broader panel of 30 cancer types, provides some alleviation to this limitation, the true predictive efficacy of AGAE score in a comprehensive pan-cancer setting awaits validation through forthcoming prospective trials involving ICIs. Secondly, there exists an incomplete record of critical clinical annotations, including but not limited to sex, age, tumor stage, TMB, and ITH, in certain RNA-Seq datasets pertaining to ICIs. This paucity of data impedes the conduct of comprehensive multivariate regression analyses and hampers efforts to comparatively assess the predictive capacities of various biomarkers. Additionally, the mechanism by which CEP55 regulates the tumor immune response requires further exploration through additional in vitro and in vivo experiments.

### Conclusions

In conclusion, the AGAE score is a promising tool for predicting clinical outcomes in cancer patients receiving ICIs. By utilizing a gene-pairing approach and advanced machine learning techniques, the AGAE score enhances predictive accuracy beyond existing biomarkers. Our findings reveal that a low AGAE score correlates with favorable immune activation, while a high score indicates immune evasion characteristics. Additionally, we identified CEP55 as a potential therapeutic target, with its knockdown improving responses to ICI therapy. The development of a user-friendly web-based tool for calculating the AGAE score provides clinicians with a valuable resource for optimizing patient selection and personalizing treatment strategies. This research highlights the need for reliable predictors of ICI response, ultimately aiming to improve treatment outcomes for cancer patients. Future validation of the AGAE score across diverse cancer types will further enhance its clinical applicability.

## Author Contributions

The study was conceived and designed by PZ, AJ, and XL. Data collection was conducted by BY. AJ performed the statistical analysis. The first draft of the manuscript was written by BY, JF, and QM. The experiment was performed by PZ and BY. The final approval of the submitted version was given by PZ, AJ, and XL. All authors contributed to the manuscript and approved the submitted version.

## Ethical statement

The procedure related to human subjects was approved by the Ethics Committee of the Changhai Hospital (No. CHEC2021091). The procedure related to animal subjects was approved by the Ethics Committee of theTianjin Medical University Cancer Institute and Hospital (No. 2023078).

## Data availability

The data used to support the findings of this study are available from the corresponding author on reasonable request.

---

**Supplementary Figure 1. A-C.** K-M survival analysis for AGAE score in the training (**A**), validating (**B**), and testing (**C**) cohorts. **D-F.** AUC values of the AGAE score in predicting 2 and 3-year survival rates.

**Supplementary Figure 2. A.** Western blot analysis of knockout of CEP55 in CT26 and 3LL cell lines.

